# A conserved mechanism for regulation of endo-lysosomal pH by histone deacetylases

**DOI:** 10.1101/252122

**Authors:** Hari Prasad, Rajini Rao

## Abstract

The pH of the endo-lysosomal system is tightly regulated by a balance of proton pump and leak mechanisms that are critical for storage, recycling, turnover and signaling functions in the cell. Dysregulation of endo-lysosomal pH has been linked to aging, amyloidogenesis, synaptic dysfunction, and various neurodegenerative disorders including Alzheimer’s disease. Therefore, understanding mechanisms that regulate luminal pH may be key to identifying new targets for treatment of these disorders. Meta-analysis of yeast microarray databases revealed that nutrient limiting conditions upregulated transcription of the endosomal Na^+^/H^+^ exchanger Nhx1 by inhibition of the histone deacetylase (HDAC) Rpd3, resulting in vacuolar alkalinization. Consistent with these findings, Rpd3 inhibition by the HDAC inhibitor and antifungal drug trichostatin A induced Nhx1 expression and vacuolar alkalinization. Bioinformatics analysis of Drosophila and mouse databases revealed that caloric control of Nhx1 orthologs DmNHE3 and NHE6 respectively, was also mediated by histone deacetylases. We show that NHE6 is a target of cAMP-response element-binding (CREB) protein, providing a molecular mechanism for nutrient and HDAC dependent regulation of endosomal pH. Control of NHE6 expression by pharmacological targeting of the CREB pathway can be used to regulate endosomal pH and restore defective amyloid Aβ clearance in an ApoE4 astrocyte model of Alzheimer’s disease. These observations from yeast, fly, mouse and cell culture models reveal an evolutionarily conserved mechanism for regulation of endosomal NHE expression by histone deacetylases and offer new therapeutic strategies for modulation of endo-lysosomal pH in fungal infection and human disease.

---

The endosome is a busy way station that handles cargo traffic at the crossroads of degradation and recycling. An important hallmark of the endo-lysosomal system is a pH gradient that is increasingly acidic through the transition from early, recycling and late endosomes to the lysosome (1). Precise tempo-spatial pH_lumen_ regulation within each of these specialized compartments is critical for a range of functions, including controlling cargo sorting, vesicular budding and formation of intraluminal vesicles, dissociation of internalized ligands, receptor recycling and turnover, modulating enzyme activity, antigen processing, and cellular signaling (2).

Endosomes are acidified by the electrogenic pumping of protons by the V-type H^+^-ATPase in conjunction with vesicular chloride transporters of the CLC family that shunt the membrane potential, effectively allowing a buildup of luminal protons (3,4). Proton leak mechanisms consisting of proton conduction channels or proton-coupled antiporters limit the acidification, and the balance of pump and leak pathways defines the pH set point. Although the molecular identity of vesicular proton channels remains poorly defined, more recently, endosomal Na^+^/H^+^ exchangers (eNHE) have emerged as the dominant leak pathway for luminal protons. Because exchanger mechanisms have high transport rates of **∼**1,500 ions/s (5), they overwhelm the capacity of the proton pump so that even small changes in eNHE activity or expression result in significant pH changes within the endo-lysosomal system.

The fundamental importance of endosomal pH is highlighted by the fact that a growing number of human diseases are associated with mutations in V-ATPase subunits (e.g., osteopetrosis, renal tubular acidosis, cutis laxa), chloride transporters (e.g., Dent’s disease) and eNHE (Christianson syndrome, autism and attention deficit hyperactivity disorder) (3-4,6). Furthermore, defective endo-lysosomal pH regulation is being increasingly linked to cellular aging, amyloidogenesis, synaptic dysfunction and neurodegenerative disorders including Alzheimer’s disease (7,8). Therefore, the discovery of novel regulators of endo-lysosomal pH may be crucial to identifying new diagnostic and therapeutic targets for these disorders.

As a starting point, we turned to the yeast *S. cerevisiae*, which has powerfully informed our understanding of fundamental mechanisms of endocytosis and vesicular traffic that are functionally conserved with higher organisms (9). By leveraging data from yeast to fly and mammalian models, we discovered an evolutionarily conserved mechanism for epigenetic control of eNHE by histone deacetylases (HDAC) in response to nutrient availability. In mammalian cells, HDAC-mediated control of endo-lysosomal pH occurs by transcriptional regulation of the endosomal Na^+^/H^+^ exchanger NHE6 (*SLC9A6*) via the cAMP-response element binding protein (CREB) pathway. We show that elevation of cAMP levels by the FDA-approved drug Rolipram results in CREB activation, increased NHE6 expression and restoration of defective Aβ clearance in the ApoE4 astrocyte model of late onset Alzheimer’s disease.

Histone acetylation has been widely recognized as a key modulator of global chromatin structure that dynamically couples extracellular signals to gene transcription. Whereas lysine acetylation on histone tails interferes with generation of higher order chromatin structures and promotes access and binding of transcription factors, histone deacetylation favors chromatin packing and represses gene transcription (10-12). Histone deacetylase (HDAC) inhibitors have been approved for the treatment of hematologic cancers and are being considered as promising therapy to reverse disease-associated epigenetic states in a range of cardiovascular, neurodegenerative and inflammatory diseases (11-14). Given their broad application, our findings offer potential therapeutic options to treat endo-lysosomal pH dysfunction in autism, AD, and other neurodegenerative disorders.

## RESULTS

### Nutrient control of endo-lysosomal pH in yeast

Yeast microarray datasets are a valuable resource for discovery-driven mining efforts. A meta-analysis of 45 microarray experiments that included 937 samples revealed increasing expression of Nhx1 with growth phase stationary>mid-log>early log (Fig. 1A), which we independently confirmed: Nhx1 transcript increased by 3.2-fold in stationary phase cells, compared to mid-log phase (Fig. 1B). Stationary phase growth in yeast is associated with depletion of nutrients. To determine if growth phase regulation of Nhx1 was linked to glucose availability, we tested the effect of glucose removal from mid-log phase yeast cultures. Nhx1 mRNA was increased by nearly 2-fold within 60 min of glucose removal (p<0.001; Fig. 1C). In contrast, addition of glucose (2%) to stationary phase cultures elicited significant decrease in Nhx1 mRNA within 30 min (p<0.001; Fig. 1D).

**Figure 1:**
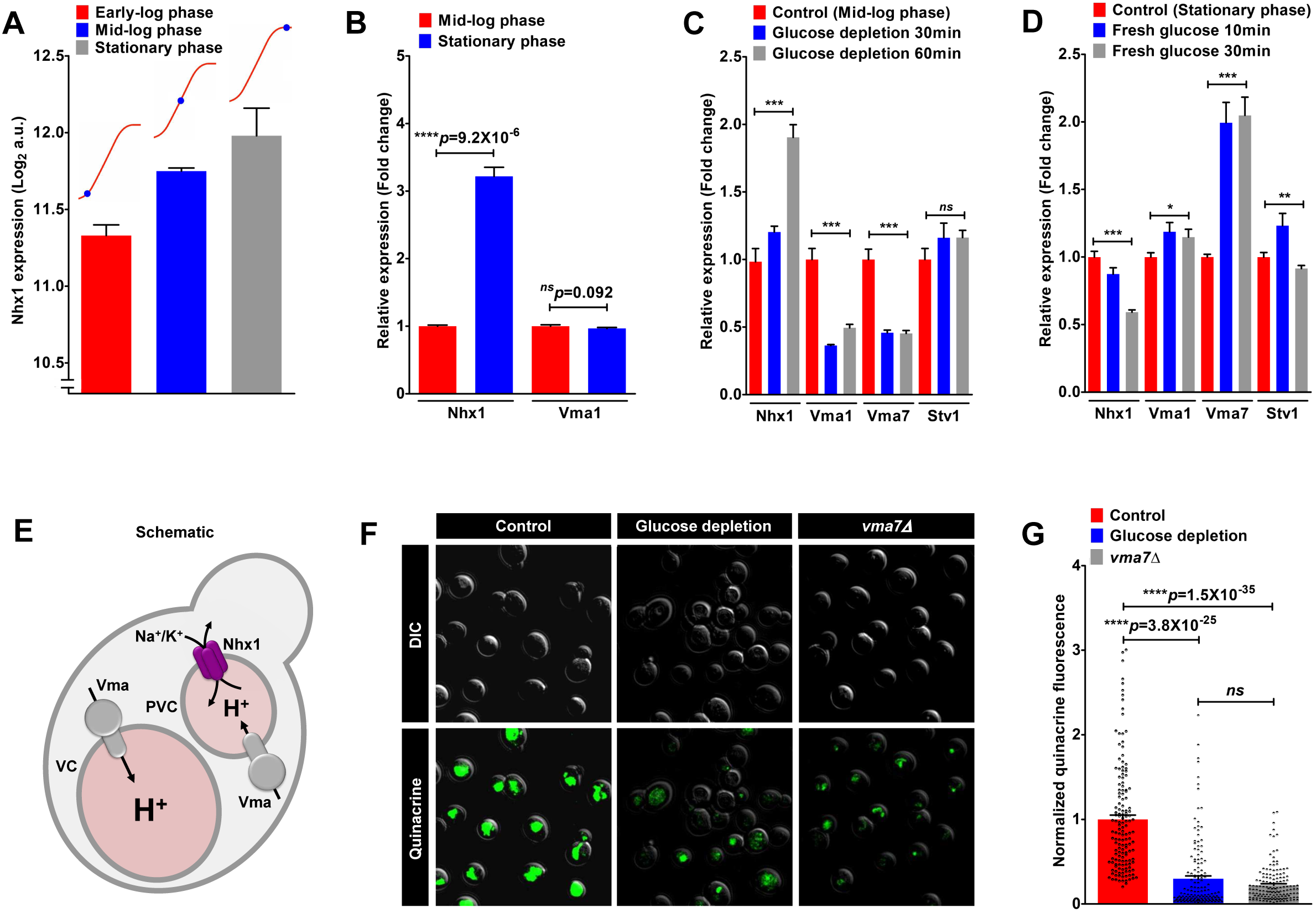
Nutrient control of endo-lysosomal pH in yeast. A. Meta-analysis of 45 microarray experiments that included 937 samples revealed increasing expression of Nhx1 with growth showing maximal expression during stationary phase (condition associated with glucose/nitrogen limitation). B. qPCR validation of Nhx1 expression showed 3.2-fold up regulation of Nhx1 transcript in stationary phase yeast, compared to mid-log phase (****p=9.2X10^-6^; *n*=3; Student’s t-test). No change in V-ATPase subunit Vma1 levels was observed (p=0.092; Student’s t-test). C. Glucose removal from mid-log phase yeast cultures showed approximately 2-fold increase in Nhx1 mRNA within 60 min of depletion (***p<0.001; *n*=3; Student’s t-test). Note significant down regulation of V-ATPase subunits with acute glucose depletion (***p<0.001, Vma1; ***p<0.001, Vma7). D. Conversely, addition of glucose (2%) to stationary phase cultures showed significant decrease in Nhx1 mRNA within 30 min (***p<0.001; *n*=3; Student’s t-test). Note significant up regulation of V-ATPase subunits with glucose addition (*p<0.05, Vma1; ***p<0.001, Vma7). (C-D) In contrast to Vma1 and Vma7, secretory pathway isoform Stv1 expression showed modest or no changes in response to glucose, suggesting that the effect of glucose regulation on the V-ATPase is largely localized to endosomal/vacuolar compartments. E. Schematic to show that Nhx1 functions as a leak pathway for protons within the endosomal/prevacuolar compartment, to balance the proton pump activity of the V-ATPase (Vma). (F-G) Representative micrographs (F) and quantification (G) showing quenching of quinacrine fluorescence with acute glucose depletion, similar to V-ATPase deficient *vma7Δ* mutants, pointing to increased vacuolar alkalization (****p<0.0001; *n*=100; Student’s t-test).

Nhx1 functions as a leak pathway for protons within the endosome/prevacuolar compartment, in opposition to the proton pump activity of the V-ATPase (15) (Fig. 1E). The balance of proton pump and leak sets the pH within the endo-lysosomal compartments. Therefore, we also analyzed expression of V-ATPase subunits under the same experimental conditions. Although we did not observe a growth phase-dependent transcriptional change in the V-ATPase catalytic subunit Vma1 (Fig. 1B), V-ATPase subunits were significantly downregulated with acute glucose depletion (p<0.001, Vma1; p<0.001, Vma7; Figure 1C) and upregulated upon glucose addition (p<0.05, Vma1; p<0.001, Vma7; Figure 1D) in contrast to Nhx1. Previous studies have shown that yeast V-ATPase subunit Stv1 is selectively targeted to the Golgi/secretory pathway and is not required for vacuolar acidification and function (16,17). In contrast to Vma1 and Vma7, Stv1 expression showed modest or no changes in response to glucose (Figure 1C-D), suggesting that the effect of glucose regulation on the V-ATPase is largely localized to late endosomal/vacuolar compartments.

We predicted that reciprocal transcriptional changes in Nhx1 and V-ATPase subunits would alter the pH of the yeast vacuole. Vacuolar pH can be probed in situ with the pH-sensitive fluorescent dye quinacrine, which accumulates in acidic environments (18). Following glucose depletion, yeast vacuoles showed little or no quinacrine staining, similar to V-ATPase deficient *vma7Δ* mutants (Fig. 1F-G; p<0.0001, n=100). Consistent with the potential role for Nhx1 in regulating nutrient control of endo-lysosomal pH in yeast, *nhx1Δ* mutants showed severely reduced survival during the stationary phase growth (19). Post-translational regulation of V-ATPase disassembly and reassembly in response to glucose concentration is well established (20,21). Here, we conclude that glucose also elicits reciprocal transcriptional changes in proton pump and leak mechanisms to regulate the pH gradient across yeast vacuoles.

### Regulation of endo-lysosomal pH in yeast is mediated by a histone deacetylase

We extended our unbiased bioinformatics approach to analyze data from 284 microarray studies that comprised a wide range of experimental conditions including gene deletion, overexpression and mutations, drug/toxin treatment, nutrient limitation or excess, metabolic/environmental stress and more. We identified experimental conditions eliciting a minimum of ±2-fold change in *NHX1* gene expression (Fig. 2A, Fig. S1A). The highest up regulation of Nhx1 was observed under conditions of (i) genetic deletion of Rpd3, (ii) glucose depletion, and (iii) nitrogen depletion. Pathway analysis revealed that loss or inhibition of histone deacetylase (HDAC) activity of Rpd3 was common to all three of these conditions (22,23)(Fig. 2B). Thus, nutrient (glucose or nitrogen) depletion elicits histone acetylation by replacement and inactivation of the HDAC/Rpd3 complex from the promoters of specific genes (22). Consistent with this finding, Rpd3 deletion phenocopies calorie restriction by enhancing lifespan of yeast cells (24,25). Conversely, all five conditions leading to maximal down regulation of Nhx1 were linked to depletion of Abf1, a transcription factor that is negatively regulated by Rpd3 (26-28) (Fig. 2A, and S1A). Recent evidence suggests that Abf1 is activated during nutritional stress (29). Consistent with this finding, Abf1 is predicted to bind to the *NHX1* promoter (Fig. S1B). For comparison, depletion of another general regulatory transcription factor Rap1 had no effect on Nhx1 expression (Fig. 2A and S1A). Thus, all top hits in Nhx1 gene expression (both up‐ and down regulation) funneled into a single pathway: the Rpd3-Abf1-Nhx1 axis (Fig. 2C).

**Figure 2:**
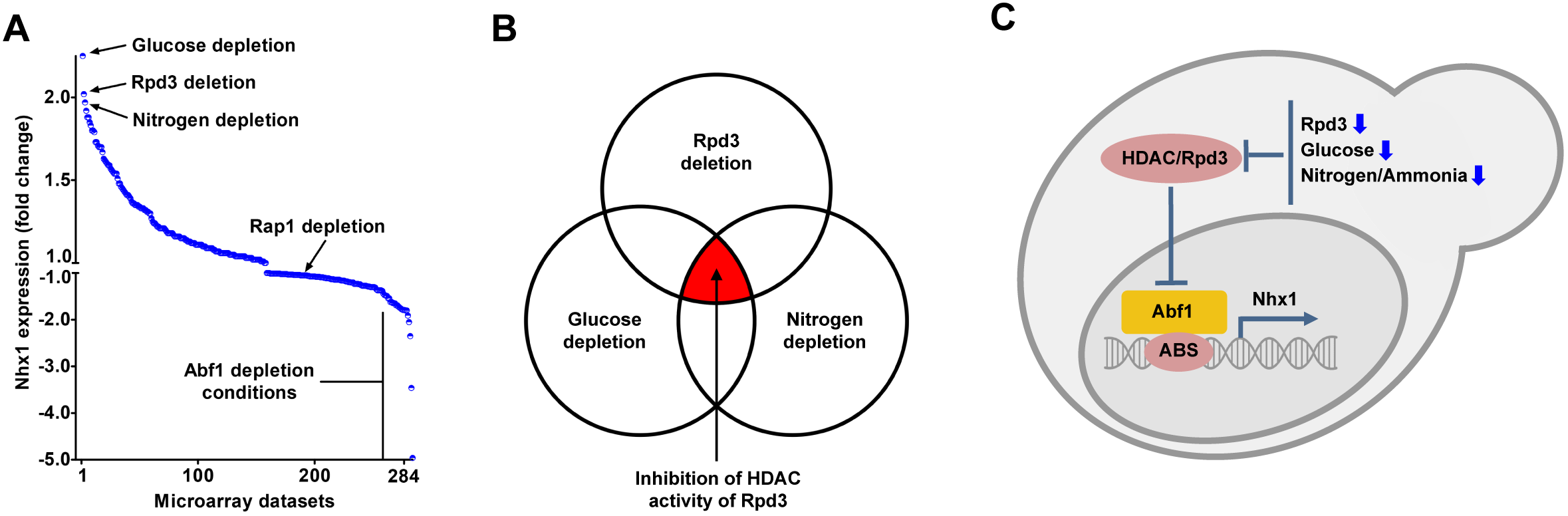
Discovery of the Rpd3-Abf1 axis for regulation of Nhx1 expression by histone deacetylases. A. Waterfall plot depicting fold-change in Nhx1 expression (y-axis) obtained from an unbiased bioinformatics analysis of 284 microarray studies (x-axis), as described in the “Experimental Methods”. Note that all five conditions leading to maximal down regulation of Nhx1 were linked to depletion of Abf1 transcription factor, known to be negatively regulated by Rpd3. B. Inhibition of histone deacetylase (HDAC) activity of Rpd3 was common to all three microarray experiments that gave maximal up regulation of Nhx1 expression, namely (i) genetic deletion of Rpd3, (ii) glucose depletion, and (iii) nitrogen depletion. C. HDAC/Rpd3-Abf1-Nhx1 axis for regulation of Nhx1 expression. This model predicts that inhibition of Rpd3 activity releases brake on Abf1 that in turn binds to Abf1 binding site (ABS) on Nhx1 promoter and enhances its expression. See related Figure S1.

To confirm the microarray data, we independently demonstrated that Nhx1 transcript was elevated by 1.9-fold in the *rpd3Δ* mutant, relative to isogenic wild type yeast (Fig. 3A). Other histone deacetylase gene deletions conferred more modest induction (1.35-fold in *hda1Δ*; Fig. 3A), or repression (0.6-fold in *sir2Δ*; Fig. 3A). All three yeast deletion strains showed similar growth (Fig. S2A). Restoration of Rpd3 by ectopic expression under a galactose-inducible promoter in *rpd3Δ* not only abolished Nhx1 upregulation in stationary phase, but also caused further repression (68% lower) in mid-logarithmic cells and inhibited yeast growth (Fig. S2B-C).

**Figure 3:**
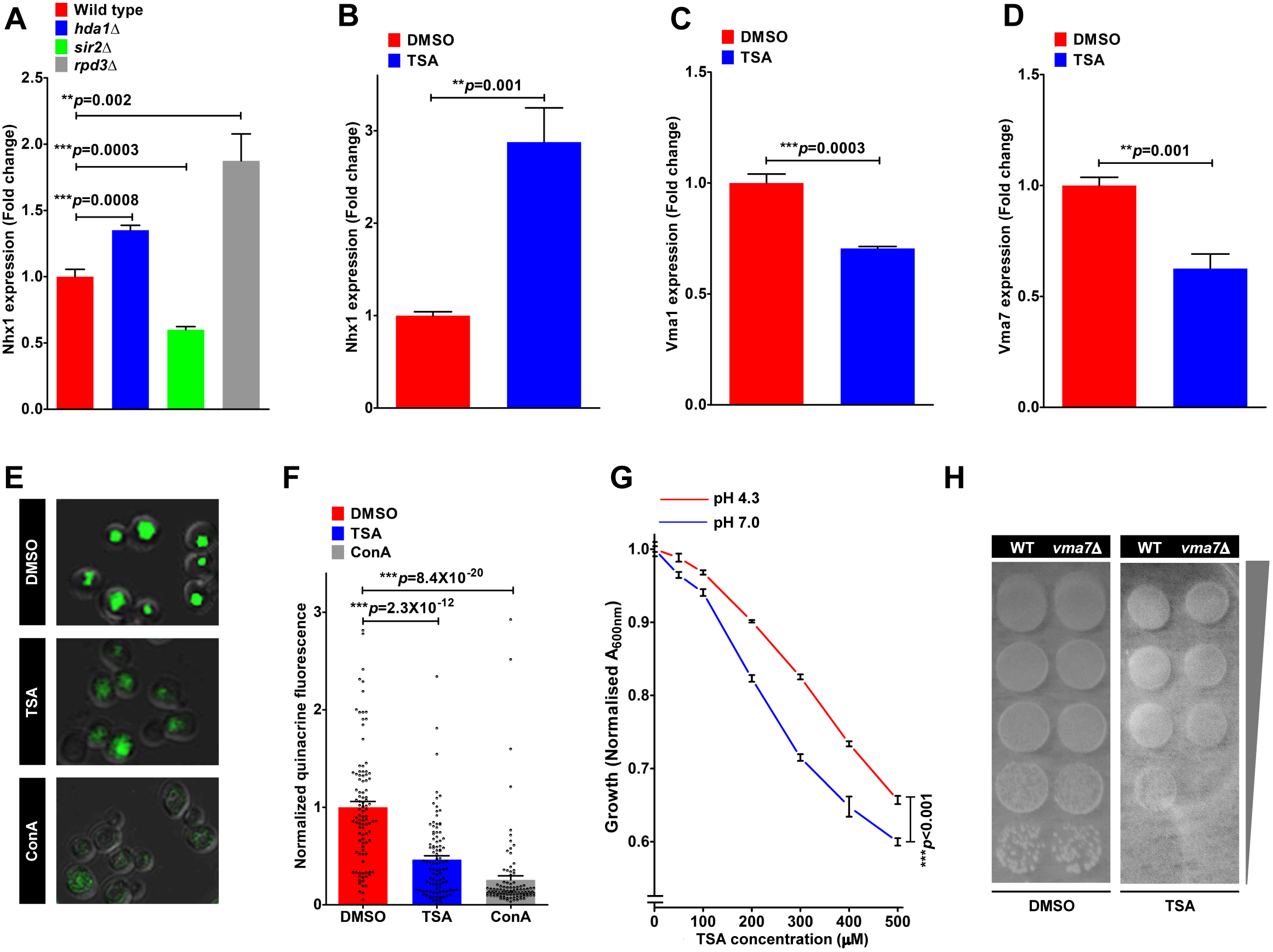
Regulation of endo-lysosomal pH in yeast is mediated by a histone deacetylase. A. Deletion of Rpd3 (*rpd3Δ* strain) increased NHX1 transcript by 1.9-fold relative to isogenic wild type yeast (**p=0.002; *n*=3; Student’s t-test). Deletion of two other histone deacetylase genes led to lesser NHX1 induction (1.35-fold in *hda1Δ*; ***p=0.0008; *n*=3; Student’s t-test), or repression (0.6-fold in *sir2Δ*; ***p=0.0003; *n*=3; Student’s t-test). B. TSA treatment of wild-type yeast elicited significant 2.88fold up regulation of Nhx1 transcript (**p=0.001; *n*=3; Student’s t-test) mimicking the effect of Rpd3 deletion. (C-D). qPCR analysis of V-ATPase subunits showed significant down regulation of (C) Vma1 (∼30% lower; ***p=0.0003; *n*=3; Student’s t-test) and (D) Vma7 (∼37% lower; **p=0.001; *n*=3; Student’s t-test) in TSA-treated yeast, compared to vehicle treatment. (E-F). Representative micrographs (E) and quantification (F) showing quenching of quinacrine fluorescence with TSA treatment, similar to treatment with a V-ATPase inhibitor concanamycin A (ConA), pointing to increased vacuolar alkalization (***p<0.0001; *n*=100; Student’s t-test). G. Alkaline pH sensitivity is a defining phenotype of increased vacuolar pH. TSA treated cells show dose-dependent reduction of growth at media pH of 7.0 as compared to pH of 4.3 (***p<0.001; *n*=4; Student’s t-test). H. Spotting assay on YPD agar plates was performed to demonstrate hypersensitivity to TSA in the *vma7Δ* yeast, lacking V-ATPase activity, indicative of synthetic lethality. See related Figure S2.

Yeast Rpd3 is a target of the histone deacetylase inhibitor and fungal antibiotic trichostatin A (TSA), and the transcriptional profile of TSA-treated wild-type yeast is similar to that of *rpd3Δ* yeast (23). We show that that TSA treatment of wild-type yeast elicited significant 2.9-fold up regulation of Nhx1 transcript (Fig. 3B) mimicking the effect of Rpd3 deletion. In contrast, we observed significant down regulation of V-ATPase subunits Vma1 (**∼**30% lower; Fig. 3C) and Vma7 (**∼**37% lower; Fig. 3D) in TSA-treated yeast, compared to vehicle treatment. In contrast to Vma1 and Vma7, the levels of Stv1 did not change significantly with TSA treatment (Fig. S2D), suggesting that the effect of TSA is specific to V-ATPase localized to the vacuolar compartment.

The transcriptional upregulation of Nhx1 and concomitant downregulation of vacuolar V-ATPase subunits by TSA is predicted to alkalinize the yeast vacuole (3). Compared to robust quinacrine fluorescence in wild-type yeast vacuoles (18), TSA treatment drastically reduced vacuolar accumulation of quinacrine, similar to the effect of the well-characterized V-ATPase inhibitor concanamycin A, pointing to increased vacuolar pH (Fig. 3E-F). Alkaline pH sensitivity is a defining phenotype of V-ATPase mutants, indicative of the reduced ability to acidify vacuoles (30,31). We found that TSA treated cells show dose-dependent reduction of growth at alkaline media pH (Fig. 3G). Hypersensitivity to TSA in the *vma7Δ* yeast, lacking V-ATPase activity, is evidenced by synthetic lethality (Fig. 3H). TSA synergizes with the antifungal action of an azole drug fluconazole that we previously showed to act via inhibition of V-ATPase activity (30)(Fig. S2E-F). Together, these data provide strong evidence for regulation of yeast vacuolar pH by HDAC inhibition.

### Evolutionary conservation of eNHE regulation by nutrient availability and histone deacetylases

To investigate if the mechanism of upregulation of eNHE in response to nutrient limitation and HDAC activity is evolutionarily conserved, we evaluated publicly available expression data in the fruit fly *Drosophila melanogaster*. Fruit fly has one eNHE isoform DmNHE3, which, like its mammalian cousin NHE6, is highly expressed in the brain (32) (Fig. 4A). Analysis of a microarray dataset studying the effects of starvation in Drosophila larvae (GSE14531) revealed significant downregulation of class I HDAC DmRpd3 (-2.14-fold; Fig. 4B) and reciprocal upregulation of class III HDAC DmSir2 (1.66-fold; Fig. 4C) in fly larvae starved for 24 h (33). Importantly, consistent with findings from yeast, there was significant upregulation (2.12-fold; Fig. 4D) of DmNHE3 expression with starvation (33). No changes in expression with starvation were seen for other NHE isoforms, DmNHE1 and DmNHE2 (Fig. S3A-B). We also evaluated the effect of nutrient depletion on transcript levels of Vha68 and Vha14 that are orthologs of Vma1 and Vma7, respectively. Unlike our observations in yeast, no change in expression of these V-ATPase subunits was noted in Drosophila, suggesting that the transcriptional effect of starvation might be limited to endosomal NHE in higher eukaryotes.

**Figure 4:**
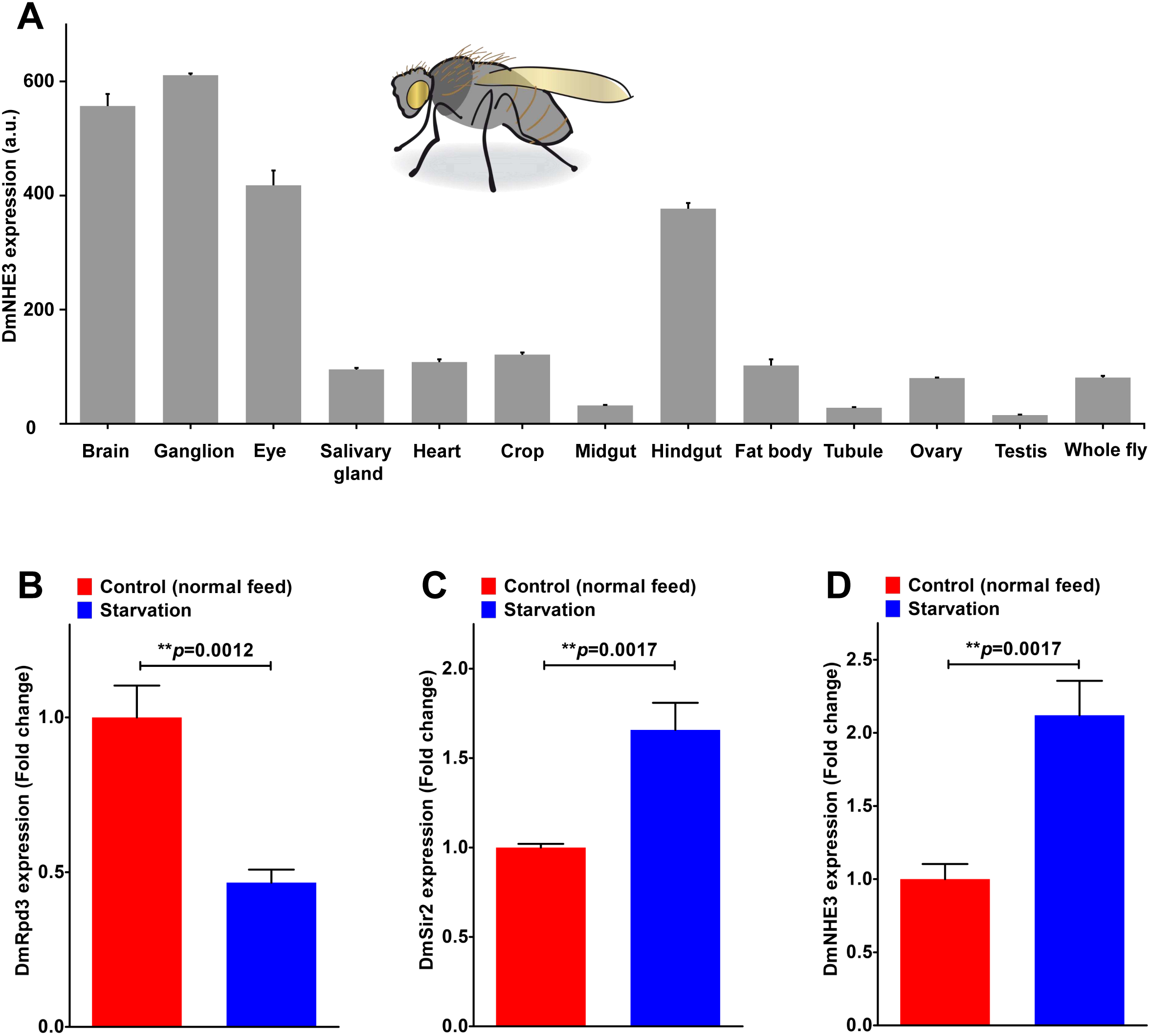
Evolutionary conservation of eNHE regulation by nutrient availability and histone deacetylases. A. Expression pattern of Drosophila eNHE isoform DmNHE3. Note high expression of DmNHE3 in the brain, like its mammalian cousin NHE6. Data obtained from FlyBase atlas. (B-C). Gene expression data showing significant down regulation of class I HDAC DmRpd3 (-2.14-fold; **p=0.0012; *n*=3; Student’s t-test; B) and reciprocal up regulation of class III HDAC DmSir2 (1.66-fold; **p=0.0017; *n*=3; Student’s t-test; C) in fly larvae starved for 24h. D. Significant up regulation (2.12-fold; **p=0.0017; *n*=3; Student’s t-test) of DmNHE3 expression with starvation suggests evolutionarily conserved mechanism for regulation of eNHE expression by histone deacetylases in yeast and fly models. See related Figure S3.

Chromatin-modifying complexes are conserved between yeast and mammals. Eighteen mammalian HDACs have been described, grouped into four functional classes based on similarity to yeast orthologs (10). Independent studies have shown that insulin, the principal hormone orchestrating response to glucose availability, alters HDAC nuclear translocation in mammalian cells (34). Calorie restriction has been shown to improve insulin sensitivity and mimic the effects caused by HDAC inhibitors, including on histone acetylation and activation of cAMP-response element-binding (CREB) protein (35-38). Intriguingly, the mammalian eNHE ortholog NHE6 is a reported insulin responsive protein and its subcellular trafficking is regulated by insulin (39). Furthermore, in neurons, loss of NHE6 has been reported to attenuate brain-derived neurotrophic factor (BDNF) signaling that is downstream of insulin and CREB cascade (40,41). Therefore, to test if calorie restriction or HDAC inhibitors enhance mammalian eNHE expression, we first analyzed publicly available expression data on caloric restriction (approximately 10% lower than *ad lib* intake) in the mouse (Fig. 5A). These data revealed significant upregulation of NHE6 in the neocortex (Fig. 5B), but not of the closely related endosomal NHE9 or V-ATPase subunit Vma1/ATP6V1A (Fig. S4A-B) (42). Next, we analyzed two independent microarray experiments studying the effect of TSA in cell culture at shorter (2 h; GSE8488) and longer (18 h; GSE9247) time points (43,44). Expression data from both these experiments revealed significant up regulation of NHE6 with TSA treatment (Fig. 5C-D), but not NHE9 (Fig. S5A-D) or NHE1 (Fig. S5B-E). Activation of transcription factor CREB is a common response to both calorie restriction and pharmacological HDAC inhibition (12,38). Thus, for comparison, we evaluated a prototypic CREB target BDNF and documented significant upregulation with calorie restriction in mice (Fig. S4C) and HDAC inhibition in cell culture (Fig. S5C-F), as predicted. These observations from yeast, fly, mouse and cell culture models support our hypothesis of an evolutionarily conserved mechanism for regulation of eNHE expression by histone deacetylases.

**Figure 5:**
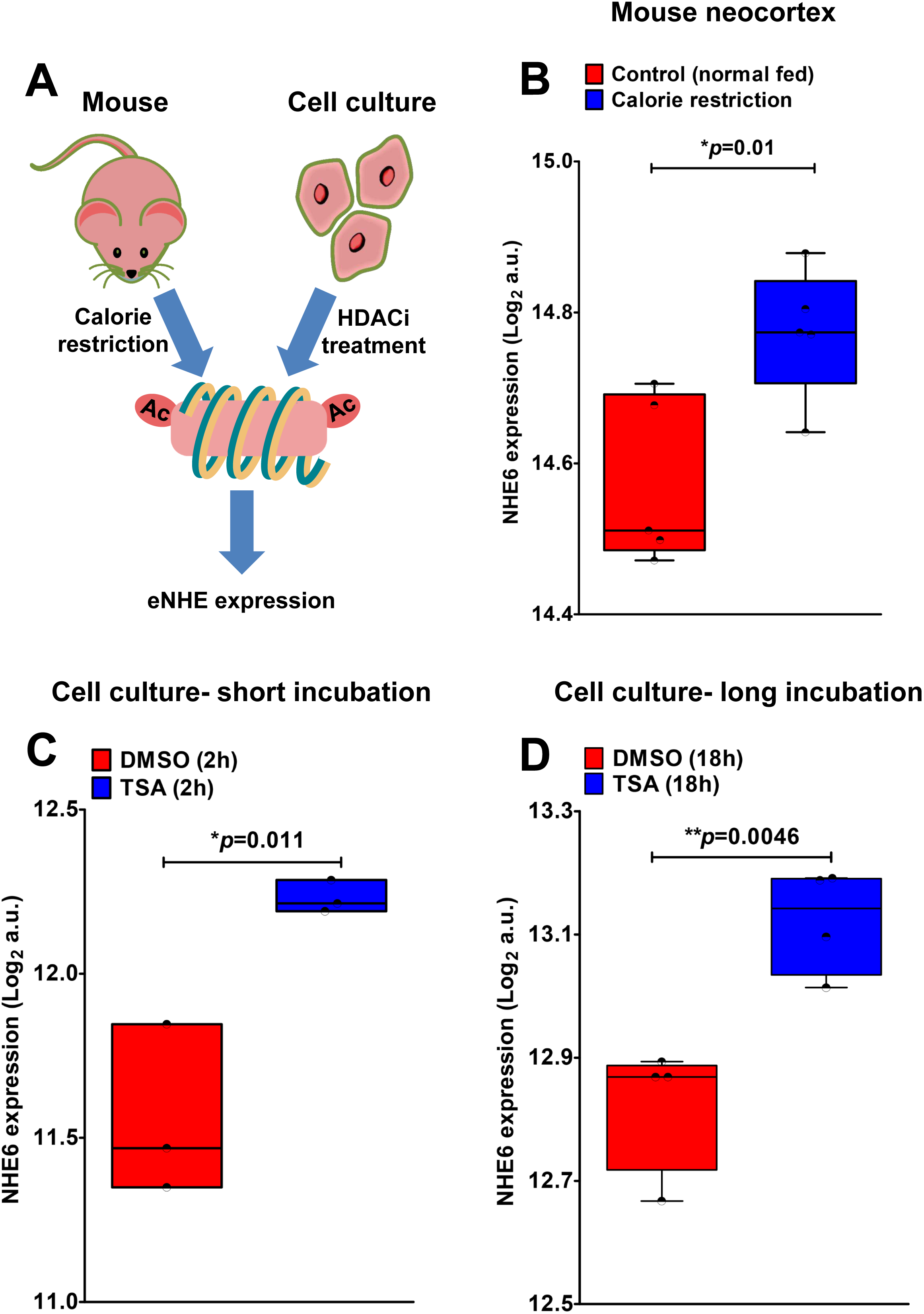
Calorie restriction and HDAC inhibition increases NHE6 expression. A. Schematic to show that dietary calorie restriction mimics the effects caused by HDAC inhibitors (HDACi) and predicted to alter eNHE expression. B. Calorie restriction (approximately 10% lower than *ad lib* intake) in the mouse significantly increased NHE6 expression in the neocortex (*p=0.01; *n*=5; Student’s t-test). (C-D). Gene expression data from two independent microarray experiments studying the effect of TSA in cell culture at shorter (2h) and longer (18h) time points. TSA treatment significantly increased NHE6 expression (p=0.011; \**n*=3; Student’s t-test; 2h; C) (**p=0.0046; *n*=4; Student’s t-test; 18h; D) at both time points. See related Figures S4 and S5.

### NHE6 is a transcriptional target of CREB

Next, we attempted to identify the HDAC-regulated transcription factor equivalent to yeast Abf1 in higher eukaryotes. In the absence of direct Abf1 orthologs in the Drosophila or human genome, we looked for parallel starvation response pathways between human and yeast. Various studies suggest that CREB transcription factor regulates response to nutrient deprivation and glucose availability in mammalian cells, analogous to the Abf1 transcription factor in yeast (38,45). Both class I (HDAC1) and class II (HDAC4) HDACs directly interact with CREB and negatively regulate its function (46,47). Furthermore, similar to the Abf1 transcription factor in yeast, the phosphorylation state of CREB is tightly regulated in response to nutrient and glucose availability (38,48). Based on all these observations, we hypothesized NHE6 as a potential CREB target (Fig. 6A).

**Figure 6:**
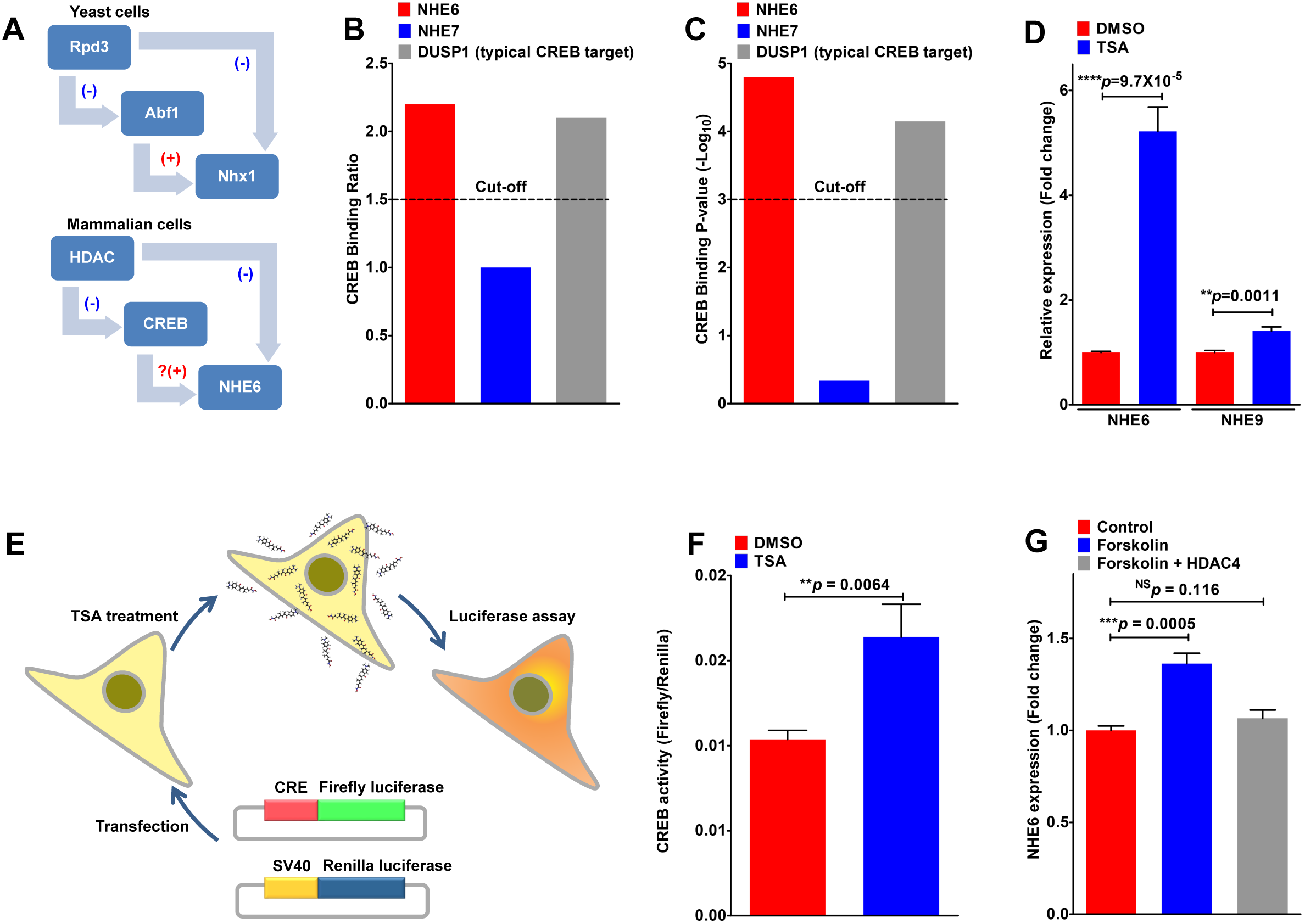
Regulation of NHE6 expression is mediated by a histone deacetylase. A. Schematic to draw parallels between regulation of eNHE between yeast (top) and mammalian cells (bottom). CREB transcription factor regulates response to nutrient deprivation and glucose availability in mammalian cells, analogous to the Abf1 transcription factor in yeast. Both class I (HDAC1) and class II (HDAC4) HDACs directly interact with CREB and negatively regulate its function. Based on this predictive model, we hypothesized NHE6 as a potential CREB target. (B-C). ChIP-on-chip (chromatin immunoprecipitation with DNA microarray) data analyzed using the recommended cutoff for ChIP positive genes (binding ratio ≥1.5 and Log_10_ p-value ≥3) revealed NHE6 as a potential CREB target. Note that the ChIP binding data for NHE6 was similar to that of DUSP1, a known CREB target, whereas no CREB occupancy was detected for the related NHE7 isoform. D. qPCR analysis demonstrating prominent ∼5.2-fold increase in NHE6 expression (****p=9.7×10^-5^; *n*=3; Student’s t-test), and modest increases in NHE9 (∼1.4-fold; **p=0.0011; *n*=3; Student’s t-test) with TSA treatment of HEK293 cells. (E-F). Luciferase assay to confirm TSA induced activation of CREB using a cyclic AMP-response element (CRE) that drives a firefly luciferase reporter gene and is measured luminometrically (E). Renilla luciferase driven by a constitutively active SV40 promoter is used to normalize for variation of cell number and transfection efficiency. Significant activation of cyclic AMP-response element (CRE) seen with TSA treatment (10μM; 12h; F) in HEK293 cells, relative to vehicle control (**p=0.0064; *n*=3; Student’s t-test). G. qPCR analysis to determine the potential of forskolin to augment the expression of NHE6 in HEK293 cells. Note significant increase in NHE6 transcript levels with forskolin treatment (***p=0.0005; *n*=3; Student’s t-test) that was blocked by expression of a constitutively active/nuclear HDAC4 mutant (S246/467/632A) (p=0.116; *n*=3; Student’s t-test). See related Figure S6.

Analysis of a publicly available dataset of CREB target genes, from chromatin immunoprecipitation with DNA microarray (ChIP-on-chip)(49), using the recommended cutoff for ChIP positive genes (binding ratio ≥1.5 and Log_10_ p-value ≥3) revealed NHE6 as a potential CREB target. The ChIP binding data for NHE6 was similar to that of DUSP1, a known CREB target, whereas no CREB occupancy was detected for the related NHE7 isoform (Fig. 6B-C). Consistent with our hypothesis in Fig. 6A, TSA treatment resulted in **∼**5.2-fold increase in NHE6 expression in HEK293 cells (Fig. 6D), whereas, modest increases were observed in NHE7 (**∼**1.3-fold; Fig S6A) and NHE9 (**∼**1.4-fold; Fig. 6D), and no changes in expression of V-ATPase subunits V0a1 (lysosomal) and V0a2 (putative endosomal) were observed (Fig. S6B-C). We used the firefly and Renilla luciferase reporter gene system to demonstrate significant activation of cyclic AMP-response element (CRE) with TSA treatment (10μM, 12 h), relative to vehicle control (p=0.0064; Fig. 6D-E). To establish a causal link between CREB activation and NHE6 expression, we treated HEK293 cells with forskolin, an adenylyl cyclase activator, and confirmed significant increase in NHE6 transcript levels that was blocked by expression of a constitutively active/nuclear HDAC4 mutant (S246/467/632A) (Fig. 6G).

To directly test if NHE6 is a CREB target, we expressed Flag-tagged CREB1 plasmid in HEK293 cells and assessed (i) functional expression of NHE6 and (ii) binding of CREB to NHE6 promoter by ChIP-qPCR (Fig. 7A). Expression of CREB1 in HEK293 cells significantly elevated NHE6 transcript (Fig. 7B), resulting in alkalization of endosomal pH from 5.9±0.03 to 6.12±0.05 (Fig. 7C). In contrast, no changes in NHE6 transcript or endosomal pH were observed with the non-phosphorylatable mutant of CREB1 (S133A; Fig. 7B-C). Chromatin immunoprecipitation was performed with a ChIP grade anti-Flag mouse antibody (Fig. 7D). Using real time qPCR we amplified three predicted CREB binding sites in the promoter of NHE6 (*SLC9A6*) at positions -12Kb, +01Kb, and +02Kb relative to transcription start site (TSS) (Fig. 7D and S6D). For comparison, we used primers targeted to NHE9 (*SLC9A9*) promoter at position -01Kb relative to TSS (Fig. 7E). We confirmed CREB binding to NHE6 promoter at all three predicted sites (Fig. 7F-H). In contrast, we observed no significant binding of CREB to NHE9 promoter (Fig. 7I). Specific pull-down of CREB1 was confirmed by probing the ChIP samples with an antibody against CREB raised in rabbit (Fig. 7J). Consistent with our ChIP-qPCR findings, we note that the pattern of NHE6 expression closely parallels that of CREB1 through normal human brain development (Fig. S7K; Pearson correlation: 0.62) and in all areas of the brain (Fig. 7L). In contrast, NHE9 expression showed no correlation with CREB (Fig. 7M; Pearson correlation: -0.018). Taken together, our findings reveal a novel mechanism of CRE mediated transcriptional regulation of NHE6 and provides a molecular mechanism for nutrient and HDAC dependent regulation of endosomal pH.

**Figure 7:**
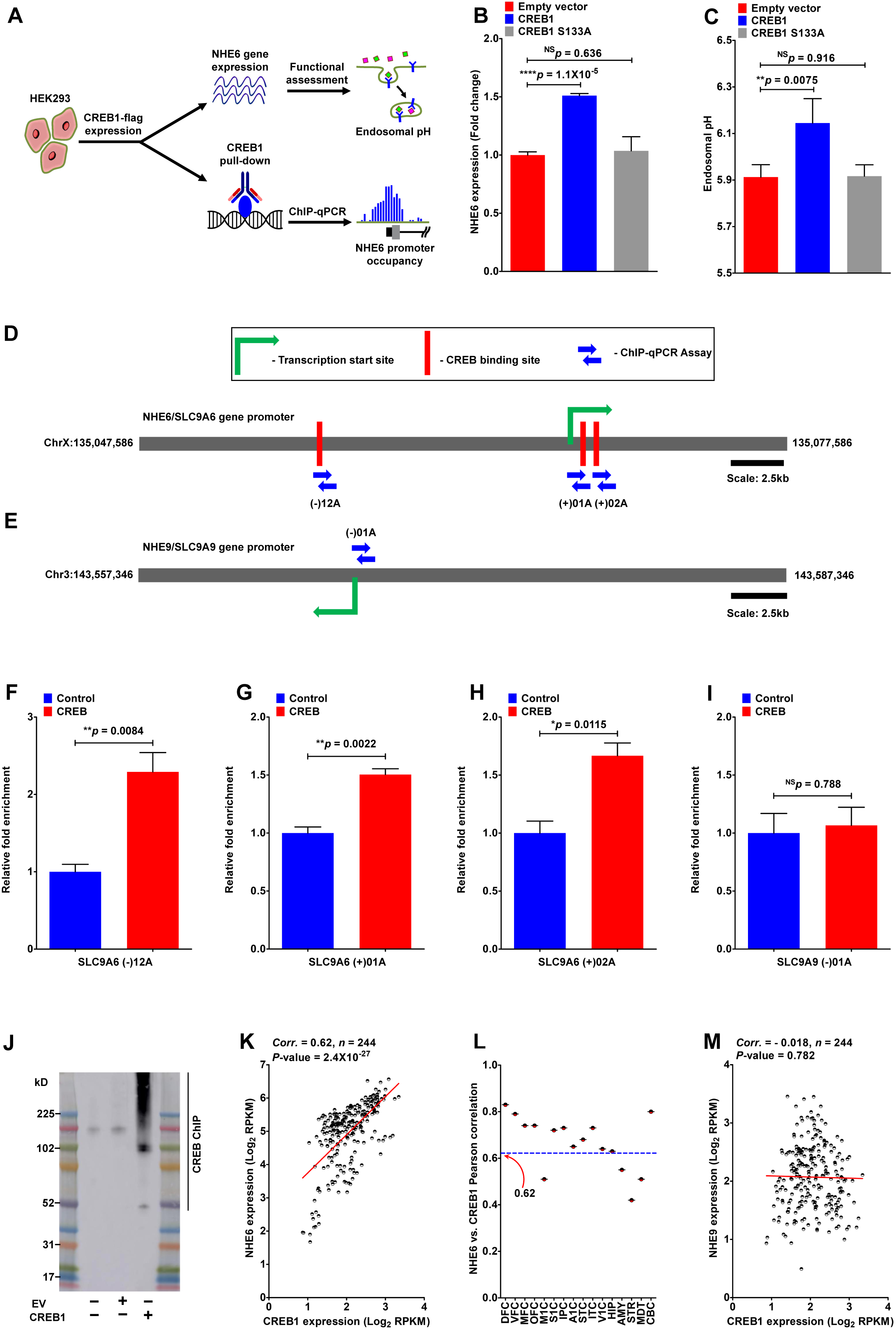
CREB mediated transcriptional regulation of NHE6 expression. A. Experimental framework to directly test if NHE6 is a CREB target. Flag tagged CREB1 plasmid was expressed in HEK293 cells and (i) NHE6 expression and its functional outcome (elevation of endosomal pH) and (ii) binding of CREB to NHE6 promoter by ChIP-qPCR were assessed. (B-C). Expression of CREB1 in HEK293 cells significantly elevated NHE6 transcript (****p=1.1X10^-5^; *n*=3; Student’s t-test; B), resulting in alkalization of endosomal pH from 5.9±0.03 to 6.12±0.05 (**p=0.0075; *n*=3; Student’s t-test; C). No changes in NHE6 transcript (p=0.636; n=3; Student’s t-test; B) or endosomal pH (p=0.916; n=3; Student’s t-test; C) were observed with the non-phosphorylatable mutant of CREB1 (S133A). (D-E). Schematic of NHE6 (top; D) and NHE9 (bottom; E) promoter element showing transcription start site, CREB binding sites, and locations of ChIP-qPCR assays. (F-H). ChIP-qPCR amplification of three predicted CREB binding sites in the promoter of NHE6 at positions -12Kb, +01Kb, and +02Kb relative to transcription start site (TSS). Note significant CREB binding to NHE6 promoter at all three predicted sites. I. No significant binding of CREB to NHE9 promoter was identified using a primer targeting position -01Kb relative to TSS. J. Western blot to confirm specific pull-down of CREB1 by probing the chromatin immunoprecipitation (ChIP) samples with an antibody against CREB raised in rabbit. ChIP was performed using an antibody raised in mouse. Note high molecular weight smear due to formaldehyde crosslinking. (K-L). Pattern of NHE6 expression closely parallels that of CREB1 through normal human brain development (K) (Pearson correlation: 0.62; *n*=244; ****p=2.4X10^-27^) and in all areas of the brain (L). Abbreviation: DFC, dorsolateral prefrontal cortex (CTX) (Brodmann area 9 (BA9), BA46); VFC, ventrolateral prefrontal CTX (BA44, BA45); MFC, medial prefrontal CTX (BA32, BA33, BA34); OFC, orbital frontal CTX (BA11); M1C, primary motor CTX (BA4); S1C, primary somatosensory CTX (BA1–BA3); IPC, posterior inferior parietal CTX (BA40); A1C, primary auditory CTX (BA41); STC, posterior superior temporal CTX (BA22); ITC, inferior temporal CTX (BA20); V1C, primary visual CTX (BA17); HIP, hippocampal formation; AMY, amygdaloid complex; STR, striatum; MDT, mediodorsal nucleus of thalamus; CBC, cerebellar CTX. M. NHE9 expression showed no correlation with CREB1 (Pearson correlation: -0.018; n=244; p=0.782). Data obtained from Allen Brain atlas. See related Figure S6.

### Control of NHE6 expression by pharmacological targeting of CREB pathway

Christianson syndrome (CS) patients, with loss of function mutations in NHE6, clinically present with neurodevelopmental disorders of intellectual disability and severe autism, and show age-dependent hallmarks of neurodegeneration, including loss of cerebellar and cortical mass, prominent glial pathology and hyper-phosphorylated tau deposition (3-50-52). Recent revelations on the shared pathology of defective Aβ clearance and altered processing of amyloid precursor protein (APP) between autism and Alzheimer disease suggest that endosomal pH regulation may be one critical mechanistic link between neurodevelopmental and neurodegenerative disorders (53). NHE6 gene expression is depressed in both AD and autism (54,55). CRE mediated transcriptional regulation of NHE6 expression offers opportunities to expand the repertoire of drugs that could potentially correct human pathologies resulting from NHE6 down regulation and aberrant endosomal hyperacidification (Fig. 8A). For example, Rolipram, a PDE4 inhibitor that elevates cAMP levels resulting in CREB activation, has shown efficacy as memory enhancer and for amelioration of Aβ related neuropathology in mice models (56).

**Figure 8:**
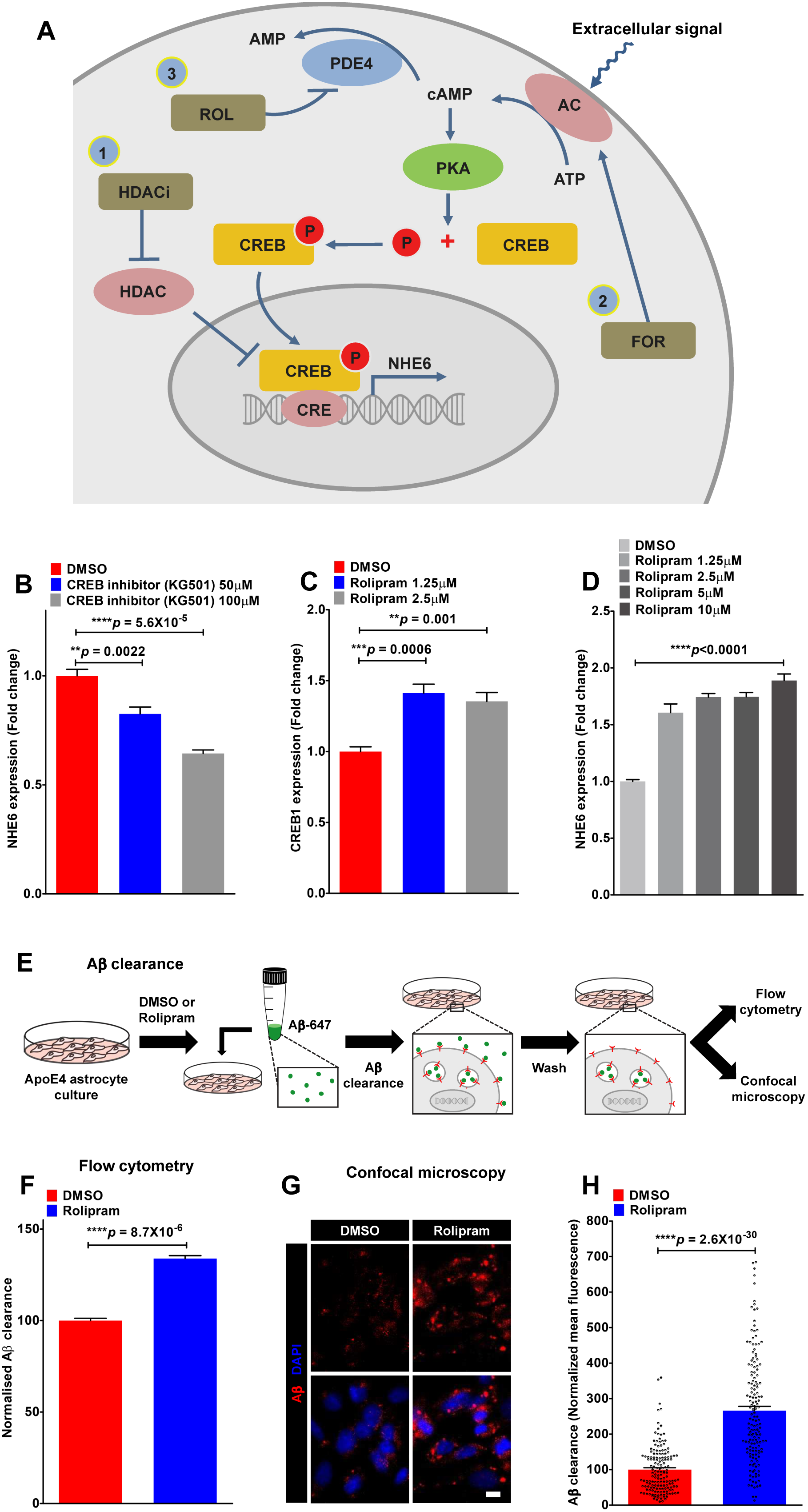
Control of NHE6 expression by pharmacological targeting of CREB pathway. A. Extracellular signals induce adenylyl cyclase that elevate cAMP levels and activate the cAMP-dependent protein kinase A (PKA), which in turn induces CREB phosphorylation at serine-133. Active phospo-CREB translocate to nucleus and activates the transcription of NHE6 that contains CRE in its promoter. At least three different pharmacological approaches could be used to activate CREB and enhance NHE6 transcription. First, HDAC inhibitors (HDACi) activate CREB by attenuating HDAC-mediated repression. Second, forskolin (FOR), an agonist of adenylyl cyclase, stimulates production of cAMP and activate CREB. Third, rolipram (ROL), a PDE4 inhibitor, inhibits hydrolysis of cAMP and activate CREB. B. qPCR analysis to confirm CREB-mediated NHE6 regulation in ApoE3 astrocytes using the CREB inhibitor (KG501). Note dose-dependent down regulation of NHE6 transcript levels with KG501 treatment (****p<0.0001; *n*=3; Student’s t-test). C. qPCR analysis demonstrating dose-dependent increase in CREB1 transcript levels with rolipram treatment in ApoE4 astrocytes. D. qPCR analysis to demonstrate dose-dependent increase in NHE6 transcript levels with rolipram treatment (1.25μM to 10μM) in ApoE4 astrocytes (****p<0.0001; *n*=3; Student’s t-test). E. Fluorescent-based assay to monitor clearance of Aβ peptides by ApoE4 astrocytes. F. Quantitation of Aβ clearance from FACS analysis of 10,000 cells in biological triplicates confirmed restoration of Aβ clearance in ApoE4 astrocytes with rolipram treatment (****8.7×10^-6^; *n*=3; Student’s t-test). (G-H). Representative micrographs (G) and quantification (H) showing ∼2.66-fold increase in cell-associated Aβ in ApoE4 astrocytes with rolipram treatment (****p=2.6×10^-30^; n=160/condition; Student’s t-test). See related Figure S7.

We used mouse astrocytes with knock-in of human ApoE alleles that alter genetic risk for Alzheimer disease (57). Recently, we showed that NHE6 expression is severely reduced in astrocytes carrying the pathological ApoE4 variant, relative to the isogenic non-risk ApoE3 allele (8). Consistent with this observation, CREB is downregulated in ApoE4 brains (58). First, we confirmed CREB-mediated NHE6 regulation in ApoE3 astrocytes using the CREB inhibitor (KG501): we observed dose-dependent down regulation of NHE6 transcript levels with KG501 treatment (Fig. 8B). Rolipram was effective in dose-dependent activation of CREB targets in ApoE4 astrocytes including the prototypic CREB target BDNF (Fig. S7A), and astrocyte specific CREB targets, ALDH1A1 and FBXO2(59) (Fig. S7B-C). We also observed significant induction of CREB mRNA by Rolipram as well (Fig. 8C), consistent with similar observations in rat hippocampus (60). Here, we demonstrate dose-dependent increase in NHE6 transcript levels with Rolipram treatment (1.25μM to 10μM) in ApoE4 astrocytes (Fig. 8D). To evaluate the efficacy of Rolipram treatment on NHE6 mediated function, we monitored Aβ clearance in cultured ApoE4 astrocytes by flow cytometry and confocal microscopy (Fig. 8E). We show that Rolipram elicits significant increase in Aβ clearance, evidenced by increase in cell-associated fluorescence by flow cytometry of ApoE4 astrocytes (Fig. 8F). Similarly, endocytosed Aβ, monitored by confocal microscopy, exhibited an average of 2.66-fold increase following treatment with Rolipram (Fig. 8G-H). Taken together, our findings show that CREB activation of NHE6 expression is translatable into correction of Aβ clearance deficits in ApoE4 astrocytes.

## DISCUSSION

### Nutrient control of endo-lysosomal pH

The discovery of endosomal Na^+^/H^+^ exchangers (eNHE) in the mid-90’s and subsequent work in a range of model organisms and mammalian systems has established the fundamental importance of these ion transporters in human health and disease (3,61). Although the contribution of the V-ATPase in nutrient control of endo-lysosomal pH is well established in yeast (20,21), the role of eNHE in this mechanism was previously unknown. Here, we leveraged public data from yeast, fly and mammalian studies to make an unbiased *in silico* prediction leading to the discovery of an evolutionarily conserved mechanism for regulation of eNHE expression by nutrients and histone deacetylases (HDACs). Previous studies have established that glucose triggers rapid, post-translational regulation of disassembly and reassembly of V-ATPase to regulate the pH gradient across yeast vacuoles (20,21). In this study, we show that glucose also elicits reciprocal transcriptional changes in Nhx1 and V-ATPase subunits to control vacuolar pH. This finding is in agreement with previous work showing severely reduced survival of Nhx1 mutants during nutrient-depleted stationary phase (19).

The regulation of endo-lysosomal pH by Nhx1 and V-ATPase (or their orthologs in higher eukaryotes) might promote cell survival during nutrient depletion or starvation by at least three potential mechanisms: (i) Vacuole acidity facilitates proton-dependent import of amino acids into the lumen through Avt1, a neutral amino acid transporter important for starvation resistance (18,62). In nutrient depleted cells, decrease in vacuolar acidification would reduce vacuolar import of amino acids and help maintain cytoplasmic amino acid homeostasis (Fig. S8A-B). (ii) Transcriptional control of V-ATPase subunits could supplement post-translational mechanisms of V-ATPase disassembly as an energy-saving mechanism to conserve ATP under nutrient limiting conditions. (iii) We have previously shown that Nhx1 regulates vesicle trafficking and lysosomal biogenesis to control internalization of plasma membrane proteins and degradation in vacuoles (15,61). Increased Nhx1 expression in glucose-depleted cells might therefore help promote endocytosis and provide resources for survival during glucose starvation (63). Intriguingly, eNHE mutants in *C. elegans* (Nhx5) and fly (DmNHE3) were found to be resistant to metformin, a biguanide drug commonly used to treat type-2 diabetes, potentially through dysregulation of autophagy (64). Given that metformin is known to induce a caloric restriction-like state (65), our findings suggest that upregulation of eNHE and alkalinization of endosomal pH may underlie response to metformin therapy.

### Antifungal drugs target endo-lysosomal pH

Pathogenic fungi have become a leading cause of morbidity and mortality in the world (66). Fungi need V-ATPase function to infect their hosts, and genetic inactivation of vacuolar specific V_o_ subunit Vph1, but not secretory pathway specific subunit Stv1, abolished virulence and invasive growth of *C. albicans,* underscoring the importance of endo-lysosomal pH as a drug target (16,17). Consistent with this finding, the antifungal activity of azoles, which inhibit biosynthesis of fungal-specific sterol, ergosterol, is mediated by reduced V-ATPase activity in ergosterol-deficient vacuoles (30). Similarly, decreased V-ATPase activity may underlie the antifungal activity of amphotericin B, which is known to directly bind and sequester ergosterol in yeast cells (67). Furthermore, many naturally occurring antifungal compounds such as terpenoid phenols, function at least in part by disrupting V-ATPase function (68). Recent studies have documented the potential benefits of repurposing FDA approved amphipathic drugs such as amiodarone, already in clinical use as calcium antagonists, that partition into acidic compartments, alkalinize endo-lysosomes and potentiates antifungal effects and reverse resistance to azole drugs (69). Here we show that an antifungal HDAC inhibitor trichostatin A (TSA) increases vacuole pH and enhances the antifungal effects of fluconazole. Intriguingly, both amiodarone and HDAC inhibitors confer significant neuroprotection in cell culture and animal models (11-12,70). Together, these studies and our current findings makes a strong case for (i) repurposing antifungal drugs as potential drugs to correct human pathologies arising from dysregulation of endolysosomal pH and (ii) repurposing FDA approved drugs having off-target effects on pH homeostasis as candidates for the development of new therapeutic strategies for fungal disease.

### Targeting endo-lysosomal pH in human diseases

Dysregulation of endo-lysosomal pH is an emerging theme in pathogenesis of a wide range of human diseases, ranging from neurodegeneration to cancer (7,71). There is clearly potential for intervention to exploit the disease-modifying effects of endosomal pH, and this should be investigated further. From lessons learnt with Baker’s yeast, we known that gene disruption of *NHX1* leads to cellular phenotypes reminiscent of Alzheimer’s disease (AD) that include enlarged endosome/prevacuolar compartment, hyperacidic luminal pH, enhanced proteolysis of the chaperone protein Vps10 (a homolog of the AD susceptibility factor SORL1), mistrafficking of vacuolar hydrolases and profound deficiencies in lysosomal cargo delivery (15,72). Several studies have identified links between eNHE and a host of neurological disorders including autism, intellectual disability, epilepsy, Parkinson’s disease, multiple sclerosis and more recently to late-onset AD (LOAD) (3-8,54,73), although mechanisms underlying eNHE regulation have remained obscure. Here we show that expression of the Christianson Syndrome protein NHE6 is under control of CREB transcription factor. A translatable finding from our studies is that pharmacological HDAC inhibition (by TSA) or CREB activation (by Forskolin or Rolipram) elevate endosomal pH and have the potential to mitigate human pathologies such as AD and autism associated with aberrant endosomal hyperacidification. Furthermore, regulation of endo-lysosomal pH may at least partially explain the working mechanism of HDAC inhibitors in cancer therapy. Tumor cells in the periphery have better blood supply, whereas cells inside tumor masses are under starved conditions, since nutrients are not fully supplied to these cells. As we have shown earlier, a subset of brain cancer glioblastoma multiforme show significant upregulation of eNHE isoform NHE9 that might help in survival of tumor cells in nutrient limited conditions (74). Together, these data suggest that, by governing fundamentally critical endosomal pH homeostasis, eNHE plays an important role in mediating nutrient control of endo-lysosomal pH. This importance makes them an attractive target as antifungal drugs and in several human diseases. Exploration of the conserved mechanism for regulation of endo-lysosomal pH by histone deacetylases provides a foundation for understanding the pathways regulating pH homeostasis and a platform for the development of drugs that could potentially correct human pathologies resulting from dysregulated endo-lysosomal acidification.

## EXPERIMENTAL PROCEDURES

### Yeast strains and growth

All *Saccharomyces cerevisiae* strains used were derivatives of BY4741(gift from Dr Susan Michaelis, Johns Hopkins University or purchased from transOMIC technologies). Yeast cells were grown in YPD (yeast extract, peptone and dextrose) medium at 30°C with shaking at 250 rpm. Acute glucose depletion was performed by growing yeast in in YP (yeast extract and peptone) medium. Yeast growth was determined by optical density of the culture at 600 nm (OD600). Spotting assay on YPD agar plates was performed to detect drug-induced growth inhibition phenotypes.

### Alkaline pH sensitivity and quinacrine staining

Increased luminal pH in the yeast vacuole was detected using two defining phenotypes: alkaline pH sensitivity and quenching of quinacrine staining. Sensitivity to alkaline media pH was conducted in YPD medium, which was adjusted to pH 7 or pH 4.3. For quinacrine staining, cells were suspended in YPD medium containing 100 mM HEPES pH 7.6, 200 μM quinacrine and stained for 10 min at 30°C. Stained cells were washed 3X with 100 mM HEPES pH 7.6, 0.2% glucose and examined by fluorescence microscopy immediately. Quantification of images was done using ImageJ software.

### Cell Culture

Human ApoE isoform-expressing (ApoE3 and ApoE4) immortalized astrocytes (gift from Dr. David M Holtzman, Washington University, St. Louis) were maintained in DME-F12 (Invitrogen) supplemented with 10% fetal bovine serum (FBS) (Invitrogen) and 200μg/ml Geneticin/G418 (Corning Cellgro). HEK293T cells were obtained from ATCC and grown in DMEM/high glucose/sodium pyruvate (Invitrogen) containing 10% FBS. Culture conditions were in a 5% CO2 incubator at 37°C. Cell viability was measured using the trypan blue exclusion method.

### Aβ clearance assays

To quantify Aβ clearance, ApoE4 astrocytes were washed with serum-free medium (SFM) followed by incubation with 100 nM fluorescently-labeled HiLyte Fluor 647-Aβ40 (æAS-64161, AnaSpec). Following several washed with PBS cells were fixed for confocal imaging using the LSM 700 Confocal microscope (Zeiss), or trypsinized for flow cytometry analysis of **∼**10,000 cells in biological triplicates using the FACSCalibur instrument (BD Biosciences). Unstained cells without any exposure to fluorescently-labeled Aβ were used as a control for background fluorescence. Quantification of confocal images was done using ImageJ software.

### Antibodies and Reagents

ChIP grade mouse monoclonal Anti-Flag (M2) (æF3165) and rabbit monoclonal Anti-CREB (48H2) (æ9197) were obtained from Sigma and Cell Signaling Technology, respectively. TSA (æA8183) was from ApexBio Technology. Concanamycin A (æC9705), Fluconazole (æF8929), KG501 (æ70485), and Rolipram (æR6520) were obtained from Sigma. Forskolin (æ3828) was purchased from Cell Signaling Technology.

### Plasmids and transfection

pBY011 Rpd3 plasmid under galactose inducible promoter was obtained from Harvard PlasmID Repository (æScCD00010538). Empty vector with uracil resistance was a gift of Dr Steven Claypool (Johns Hopkins University). Yeast transformation was performed using lithium acetate method. pCF CREB1 (Addgene plasmid æ22968) and pCF CREB1 S133A (Addgene plasmid æ22969) were a gift from Dr Marc Montminy. pcDNA-HDAC4 (S246/467/632A) (Addgene plasmid æ30486) was a gift from Dr Tso-Pang Yao. HEK293T cells were transfected using Lipofectamine 2000 reagent, as per the manufacturer’s instructions.

### Quantitative Real-time PCR

mRNA was extracted from yeast and cell cultures using the RNeasy Mini kit (æ74104, Qiagen) with DNase I (æ10104159001, Roche) treatment, following the manufacturer’s instructions. Yeast lysis buffer contained 1 M sorbitol, 0.1 M EDTA, 0.1% β‐ME, and zymolase (æE1004, Zymo Research). Complementary DNA was synthesized using the high-Capacity RNA-to-cDNA Kit (æ4387406, Applied Biosystems). Quantitative real-time PCR analysis was performed using the 7500 Real-Time PCR System (Applied Biosystems) using Taqman Fast universal PCR Master Mix (æ4352042, Applied Biosystems). Taqman gene expression assay probes used in this study are: Yeast: ACT1, Sc04120488_s1; NHX1, Sc04115315_s1; RPD3, Sc04161394_s1; STV1, Sc04153492_s1; VMA1, Sc04108170_s1; VMA7, Sc04124849_s1.Human:ATP6V0A1, Hs00989334_m1; ATP6V0A2, Hs00429389_m1; GAPDH,Hs02786624_g1;NHE6, Hs00234723_m1; NHE7, Hs01078624_m1; NHE9, Hs00543518_m1. Mouse: ALDH1A1, Mm00657317_m1; BDNF, Mm04230607_s1; CREB1,Mm00501607_m1;FBXO2, Mm00805188_g1; GAPDH, Mm99999915_g1; NHE6, Mm00555445_m1. The Ct (cycle threshold) values were first normalized to endogenous control levels by calculating the ΔCt for each sample. These values were then analyzed relative to control, to generate a ΔΔCt value. Fold change was obtained using the equation, expression fold change=2^-ΔΔCt^. Each experiment was repeated three times independently.

### Endosomal pH measurement

Endosomal pH was measured using flow cytometry, as we previously described (54,73). Briefly, cells were rinsed and incubated in SFM for 30min, to remove residual transferrin and then incubated with pH-sensitive FITC-Transferrin (æT2871, Thermo Fisher Scientific) (75 μg/ml) together with pH non-sensitive Alexafluor 633-Transferrin (æT23362, Thermo Fisher Scientific) (25 μg/ml) at 37°C for 1hr. Cells were placed on ice to stop transferrin uptake and excess transferrin was removed by washing 2X with ice-cold serum-free DMEM and PBS. Bound transferrin was removed by washing with ice-cold pH 5.0 PBS and pH 7.0 PBS. Cells were trypsinized and pH was determined by flow cytometry analysis of **∼**10,000 cells in biological triplicates using the FACSCalibur instrument (BD Biosciences). A four-point calibration curve with different pH values (4.5, 5.5, 6.5 and 7.5) was generated using Intracellular pH Calibration Buffer Kit (æP35379, Thermo Fisher Scientific) in the presence of 10μM K^+^/H^+^ ionophore nigericin and 10μM K^+^ ionophore valinomycin.

### Bioinformatics

Unbiased *in silico* approach was used to identify experimental conditions that significantly altered Nhx1 expression, by analyzing data from 284 Saccharomyces cerevisiae microarray studies, that comprised a wide range of studies including gene deletion, overexpression and mutations, drug/toxin treatment, nutrient limitation or excess, metabolic/environmental stress and many others. Experimental conditions with a minimum gene expression fold change of ±2 were selected for further thorough pathway analysis. A predictive model for regulation of Nhx1 expression was created based on the overlap of pathways from experimental conditions that resulted in highest degree of changes in gene expression (both up-and downregulation) and selected for *in vitro* validation. Meta-analysis of 45 microarray experiments including 937 samples determined effect of growth phase on Nhx1 expression. Normalized yeast expression data was obtained from Genevestigator (Nebion AG). Drosophila microarray experiment GSE14531 was analyzed to study starvation effect on fly larvae. Mammalian gene expression datasets included in the study were GSE11291, GSE8488 and GSE9247. Human brain and Drosophila gene expression data were obtained from the Allen Brain and FlyBase atlas, respectively.

### Chromatin Immunoprecipitation (ChIP)-qPCR

To assess physical binding of the flag tagged CREB to the NHE6 Gene promoter, ChIP-qPCR was carried out. ChIP was performed using the EpiQuik Chromatin Immunoprecipitation (ChIP) Kit (æP-2002, Epigentek), according to the manufacturer’s instructions. In brief, cells were treated with 1% formaldehyde for 10 minutes to cross-link proteins to DNA. To generate DNA fragments approximately 200 to 1000 bp long, the cross-linked chromatin was sonicated on ice. ChIP assays were performed using using a ChIP-grade Anti-flag antibody. Specific pull-down of CREB1 was confirmed by probing the ChIP samples with an antibody against CREB raised in rabbit using Western blotting. After reversal of the cross-links, qPCR was performed using the EpiTect ChIP-qPCR primer assay for NHE6/*SLC9A6* (GPH1013950(-)12A, ‒12 kb from the transcription start site; GPH1013950(+)01A, +1 kb from the transcription start site; and GPH1013950(+)02A, +2 kb from the transcription start site). ChIP-qPCR primer assay for NHE9/*SLC9A9* promoter (GPH1023313(-)01A, -1 kb from the transcription start site) was used as a control (all Qiagen). The resulting amplification and melt curves were analyzed to ensure specific PCR product. Threshold cycle (Ct) values were used to calculate the fold enhancement.

### Luciferase assay

pCRE-Luc (encoding for the cAMP-responsive element CRE and firefly luciferase), and pSV40-RL (encoding for a SV40 promoter and Renilla luciferase) plasmids (gift of Dr Jennifer L. Pluznick, Johns Hopkins University) were transiently transfected into HEK293T cells using lipofectamine 2000 reagent, as per manufacturer’s instructions. The ratio of firefly luciferase and Renilla luciferase was measured using the Dual-Luciferase Assay System (æE1910, Promega) with data collected using a FLUOstar Omega automated plate reader (BMG LabTech) to evaluate the CREB activation following treatment with a HDAC inhibitor.

## Acknowledgements

We thank Dr. David M Holtzman, Washington University, St. Louis for the gift of ApoE immortalized astrocytes. We thank Dr. Susan Michaelis, Dr. Steven Claypool and Dr. Jennifer L. Pluznick (Johns Hopkins University) for yeast strains and plasmids. This work was made possible by support from the Johns Hopkins Medicine Discovery Fund to R.R. Additional support came from a grant to R.R. from the National Institutes of Health (DK054214). H.P. is a Fulbright Fellow supported by the International Fulbright Science and Technology Award.

## Author Contributions

H.P. designed, conducted and analyzed experiments and wrote the paper. R.R. designed and interpreted experiments and wrote the paper.

